# A metastasis-associated Pannexin1 mutant (Panx1^1-89^) forms a minimalist ATP release channel

**DOI:** 10.1101/2024.03.12.584732

**Authors:** Junjie Wang, Carsten Mim, Gerhard Dahll, Rene Barro-Soria

**Author notes:** Correspondence: Rene Barro-Soria Department of Medicine, Miller School of Medicine, University of Miami, 1600 NW 10th Avenue, Miami, FL 33136, USA. Gerhard Dahl. Department of Physiology & Biophysics Miller School of Medicine, University of Miami, 1600 NW 10th Avenue, Miami, FL 33136, USA.

## Abstract

A truncated form of the ATP release channel pannexin 1 (Panx1), Panx1^1–89^, is enriched in metastatic breast cancer cells and has been proposed to mediate metastatic cell survival by increasing ATP release through mechanosensitive Panx1 channels. However, whether Panx1^1–89^ on its own (without the presence of wtPanx1) mediates ATP release has not been tested. Here, we show that Panx1^1–89^ by itself can form a constitutively active membrane channel, capable of releasing ATP even in the absence of wild type Panx1. Our biophysical characterization reveals that most basic structure-function features of the channel pore are conserved in the truncated Panx1^1–89^ peptide. Thus, augmenting extracellular potassium ion concentrations enhances Panx1^1–89^-mediated conductance. Moreover, despite the severe truncation, Panx1^1–89^ retains the sensitivity to most of wtPanx1 channel inhibitors and can thus be targeted. Therefore, Panx1 blockers have the potential to be of therapeutic value to combat metastatic cell survival. Our study not only elucidates a mechanism for ATP release from cancer cells, but it also supports that the Panx1^1–89^ mutant should facilitate structure-function analysis of Panx1 channels.

## Introduction

Extracellular ATP and its catabolites generated by ecto-ATPases are critical players in cancer biology ^1^ ^2^ ^3^ ^4^ ^5^. Thus, as an ATP release channel, Pannexin1 (Panx1) has thus gained considerable attention as a mediator of metastasis ^6–13^. Based on the direct correlation of expression of a truncated form of Panx1 channel (Panx1^1–89^) with metastatic potential of breast cancer cells, a mechanism for metastatic cell survival in the microvasculature has been proposed^14^. A key aspect of this mechanism is that the release of ATP augmented by the expression of the Panx1^1–89^ peptide. Indeed, experimental evidence indicates that released ATP facilitates the transit of the metastatic cells through the vessel wall and protects them from mechanical stress endured during transit ^14^. However, given that the mutant Panx1 truncates the protein from 426 to the amino terminal 89 amino acids, it is still unclear how the mutant Panx1 (alone or in combination with wild type Panx1) mediates ATP release.

While it is generally accepted that wtPanx1 forms an ATP release channel, presently available structural data do not support this function ^15, 16^. In all published structures of wt Panx1 or its caspase cleavage product, a constriction with a radius of 4-4.5 Å, is prominent at the extracellular entry to the channel pore ^17–23^. Since ATP has an Einstein-Stokes radius of 7 Å, it should therefore be excluded from passage through the Pannexin 1 channel. Instead, the 4-4.5 Å radius is consistent with the chloride selective conformation of the Panx1 channel. This apparent contradiction can be explained by the two-channel hypothesis ^24^. Depending on the stimulus modality, the Panx1 channel can adopt two distinct conformations ^25^: while the voltage activated or caspase activated channel is highly selective for chloride ions, a series of physiological and pathological stimuli induce the ATP permeable, large pore channel conformation ^24–29^. However, the structure of the latter conformation remains to be resolved.

In this study, we show that Panx1^1–89^ by itself can form a constitutively active membrane channel, capable of releasing ATP even in the absence of wtPanx1. Since Panx1^1–89^ forms a constitutively open channel with a pore dimension that allows ATP conduction, structural studies on Panx1^1–89^ will enhance the chance of capturing the large pore conformation of the Panx1 channel, which to date has remained elusive in all 8 published cryo-EM structures of wtPanx1. Furthermore, our biophysical characterization shows that most basic structure-function features of the channel pore are conserved in the Panx1^1–89^ peptide. Despite the severe truncation, Panx1^1–89^ retains most pharmacological properties of the wtPanx1 channel and can thus be targeted. Panx1 blockers have thus the potential to be of therapeutic value to combat metastatic cell survival.

## RESULTS

### Exclusive expression of Panx1^1–89^ results in cell death

The Panx1 mutant Panx1^1–89^ is co-expressed in metastatic cells with wtPanx1. It has been suggested that the two proteins interact, probably by assembling into hetero-oligomers, which mediate constitutive ATP release ^14^. To assess the biophysical properties of such hetero-oligomers, *Xenopus* oocytes were co-injected with mRNA for mutant and wtPanx1 proteins at a 1:1 mRNA ratio. As controls, oocytes were injected with either mutant or wtPanx1 alone.

Oocytes expressing wtPanx1 alone or co-expressing mutant and wt Panx1 survived the 24-48 hour incubation period after mRNA injection required to observe robust Panx1 channel currents. In contrast, oocytes expressing Panx1^1–89^ alone typically did not survive longer than 12 hours after mRNA injection (Figure 1a, Figure S1). These oocytes could not be voltage clamped and their appearance was distinctly abnormal. Their pigmentation at the animal pole was spotted and the cell surface exhibited indentations.

**Figure 1.**
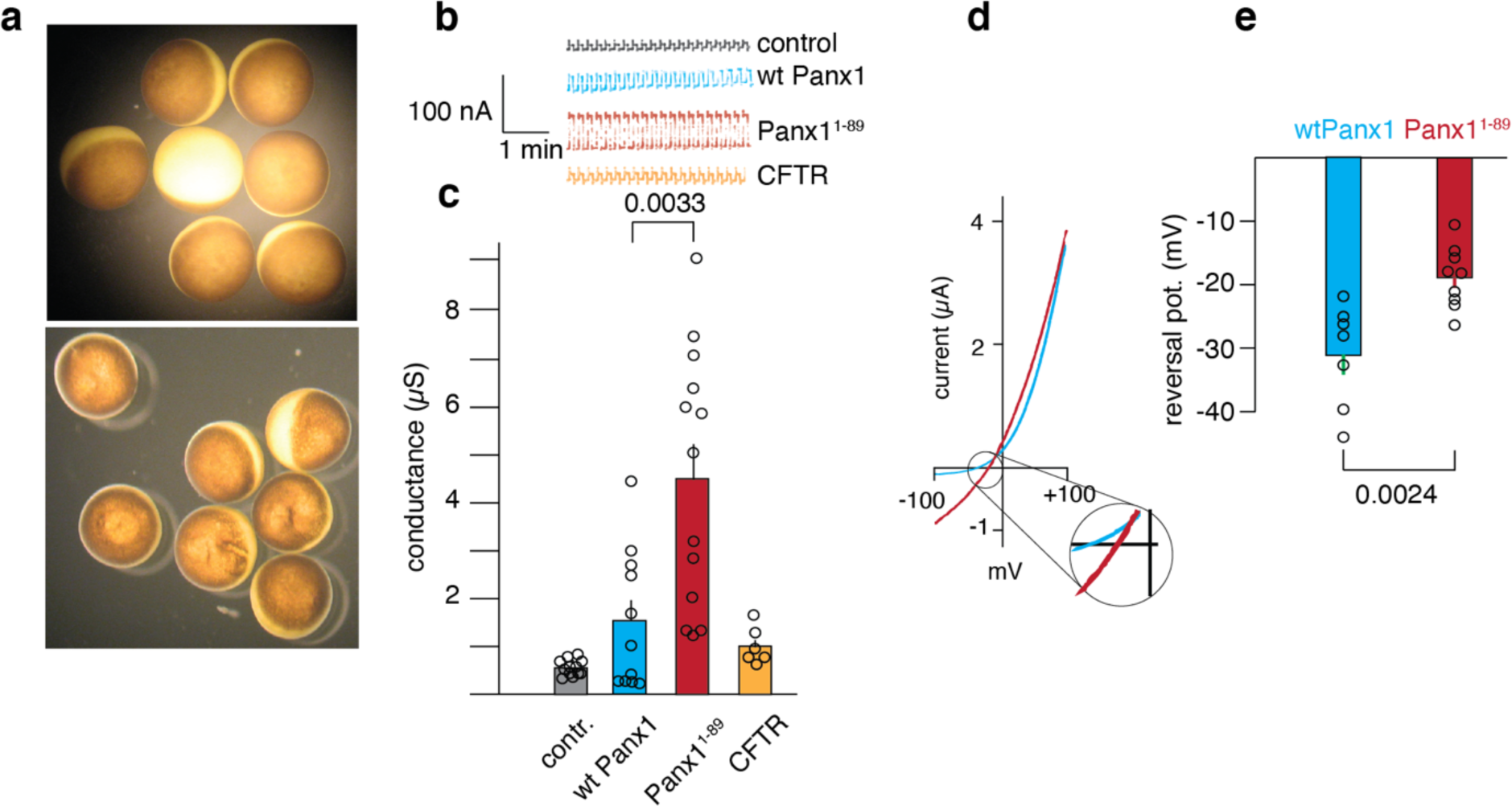
Expression of the Panx1^1–89^ mutant in *Xenopus* oocytes induces constitutive membrane currents. **a**) Control oocytes (uninjected) exhibited even pigmentation at the animal pole, no sign of swelling or shrinkage and no leakage of cytoplasmic content, indicating a healthy condition (top image). Oocytes injected with Panx1^1–89^ mRNA 24 hours prior (bottom image) exhibited uneven pigmentation, dents at the cell surface and white speckles. Such cells could not be voltage clamped, indicating total membrane breakdown. **b)** Representative current traces from uninjected (control), wtPanx1, Panx1^1–89^ and CFTR expressing oocytes, which were held at -60 mV with depolarizing 12 mV pulses. **c)** Quantitative analysis of the membrane conductance induced by the voltage steps. **d)** Voltage ramp induced current traces in oocytes expressing wt Panx1 (green) or Panx1^1–89^ (red). While the currents carried by wtPanx1 exhibited the typical outward rectification, the currents mediated by Panx1^1–89^ were prominent throughout the voltage range from -100 to +100 mV. The inset shows the area where the currents reversed from inward to outward currents. The Panx1^1–89^ mediated currents reversed at a more positive potential than the wtPanx1 currents. **e)** Quantitative analysis of the reversal potentials, which were determined on 7 (wtPanx1) and 9 (Panx1^1–89^) oocytes. Current records were obtained from oocytes 3-5 hours after injection of Panx1^1–89^ mRNA and 24-48 hours after injection of wtPanx1 mRNA. Data are presented as mean ± SEM. **b**) n=12 (contr.), 11 (wt P), 13 (P^1–89^), and 6 CFTR).

This indicated that the expression of the Panx1^1–89^ peptide alone was toxic to the cells. This toxicity could have occurred at different cellular sites, such as the synthetic pathway for proteins or at the cell membrane. In the latter case, the peptide might have created by itself a nonspecific membrane leak or could have associated with endogenous membrane proteins (channel or transporter) to confer cell toxicity. Alternatively, the peptide may have assembled in the membrane to form a channel with some resemblance to the wtPanx1 channel. The Panx1^1–89^ peptide contains only one of the four transmembrane helices of wt Panx1. Although assembly of ion channels by proteins/peptides with only one transmembrane segment is unusual, there is precedence for it. Several viroporins are oligomerized single-pass membrane peptides of a similar size range as Panx1^1–89^. Various viroporin peptides oligomerize to tetramers, pentamers or hexamers to form patent channels ^30–33^. It should be noted, however, that not all viroporins may exert a membrane channel function. For example, the SARS-Covid protein Orf3a has been shown to be involved in membrane trafficking rather than forming a channel as previously thought ^34^.

Studies using electrophysiology in combination with site-directed mutagenesis and cysteine or alanine scans have shown that the first 89 amino acid residues within the wtPanx1 channel contains key elements of the channel, including pore lining amino acids and binding sites for several modulators of channel activity (Figure S2). For example, structural and functional studies showed that amino acids W74 and R75 are involved in channel inhibition by ATP and carbenoxolone, as well as for channel activation by extracellular potassium ions ([K^+^]_o_) ^17, 35–38^.

Many of these findings were subsequently verified by determination of the Panx1 structure by cryo-EM ^17–23^. Considering the precedence given by some viroporins forming channels with a single trans-membrane segment and the presence of key features of the wtPanx1 channel in the Panx1^1–89^ sequence, it was imperative to characterize the biophysical properties of cells expressing the Panx1^1–89^ peptide heterologously.

### Expression of Panx1^1–89^ induces constitutively active membrane channels

We used two-electrode voltage clamp to measure current from Panx1^1–89^ expressing oocytes (Figure 1b, c). Compared to uninjected oocytes or oocytes expressing the chloride channel CFTR, Panx1^1–89^ expressing cells produced a robust current as early as 3 hours after mRNA injection . This is in contrast to wtPanx1, which required >12 hour incubation for currents to become detectable. Consistent with their outward rectifying properties, wtPanx1 expressing oocytes exhibited moderately increased currents when clamped at -60 mV over uninjected oocytes. Application of a voltage ramp to oocytes expressing wtPanx1 (green line in Figure 1d) showed the typical pronounced outward rectification of wtPanx1. In contrast, as shown by the large inward current at negative potentials, the channels induced by Panx1^1–89^ rectified less (red line in Figure 1d). In this respect, the Panx1^1–89^ channels are similar to the wtPanx1 channels activated by caspase cleavage at position 378 ^39^. However, while the currents mediated by wtPanx1 cleaved by caspase or truncated at position 378 reverse at the same potential as voltage activated wtPanx1 does ^16, 25, 40^, the Panx1^1–89^ induced channels exhibited a reversal potential shifted to a more positive potential (inset Figure 1d). Figure 1e shows that this shift was significant and indicates a different permeability for Panx1^1–89^-induced channels than the Cl^-^ selective property of wt and caspase cleaved Panx1 channels.

### The Panx1^1–89^-mediated current is sensitive to wtPanx1 channel blockers

Panx1^1–89^ contains amino acids within the binding sites for various Panx1 blockers ^17, 35–38^. To test whether these elements were sufficient to affect the currents mediated by Panx1^1–89^ expression, we applied several wtPanx1 blockers. Figure 2 shows that 1 mM probenecid greatly attenuated the currents although it did not completely inhibit the currents as it does in wtPanx1 channels ^41^. Other inhibitors of wtPanx1 currents, when applied in excess of their IC_50_, also attenuated the currents in oocytes expressing Panx1^1–89^, indicating a similar pharmacology for both wtPanx1 and Panx1^1–89^. Figures 2b and S3 show the effect of carbenoxolone (100µM), the food dye BB FCF (10 µM) and BzATP (100 µM). As in the case of probenecid, these Panx1 inhibitors affected Panx1^1–89^ to a lesser extent than wtPanx1.

**Figure 2.**
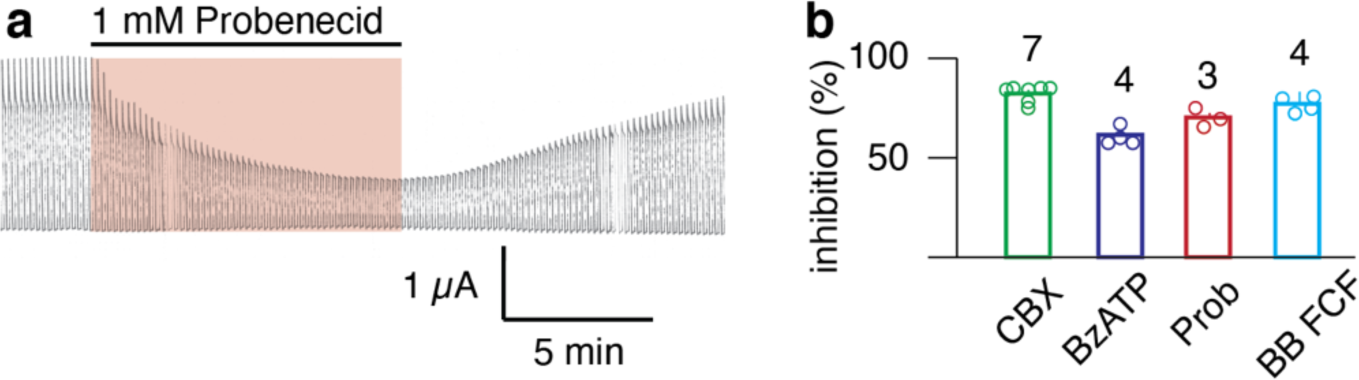
Effect of various inhibitors of Panx1 channels on the membrane currents in oocytes injected with mRNA for Panx1^1–89^. **a**) Current traces induced by repetitive voltage steps at 0.1 Hz from -60 to +60 mV. 1 mM probenecid, applied for the time indicated reversibly attenuated the currents. **b**) Quantitative analysis of % inhibition of Panx1^1–89^ -induced currents by carbenoxolone (CBX, 100 µM), BzATP (10 µM), probenecid (1mM) and brilliant blue for coloring food (BB FCF, 10 µM). Data are presented as mean ± SEM. n is indicated above the bars.

### Panx1^1–89^ responds stronger to increased extracellular potassium ion concentration than wtPanx1

The Panx1^1–89^ peptide not only contains binding sites for Panx1 blockers but also amino acids involved in the activation of the Panx1 channel at negative potentials by extracellular K^+ 37^. To test whether K^+^-activation was still retained by the truncated protein, oocytes clamped at -60 mV were perfused with a solution containing 85 mM KCl. Because Ca^2+^ attenuates the K^+^-activation of wtPanx1 ^37^, no calcium was added. Figure 3 shows the responses of uninjected oocytes and wtPanx1 or Panx1^1–89^ expressing oocytes. As reported previously, uninjected oocytes respond to extracellular high [K^+^] with a small inward current, which is significantly increased in wtPanx1 expressing cells. In contrast, the response to increased [K^+^] by Panx1^1–89^ expressing cells was on average >5 times larger. For both wtPanx1 and Panx1^1–89^, a linear relationship was observed between K^+^- induced currents and voltage induced currents. The latter was a result of voltage steps from -60 to + 60 mV and served as a measure of the expression level of the channel proteins. The slope was about 5 times steeper for Panx1^1–89^ than for wtPanx1. The linear correlation for both the wtPanx1 or Panx1^1–89^ channels argues for that K^+^-induced currents are generated by wtPanx1 and Panx1^1–89^ channels rather than an endogenous channel.

### Panx1^1-89^ expression yields a permeation pathway with large pore properties

The observation that the channels appearing in response to injection of mRNA for Panx1^1–89^ had a different reversal potential than the channels formed by wtPanx1 (Figure 1d, e) suggested that the two channels differed in their ion permeabilities. Since wtPanx1 can assume different conformations, one with chloride selectivity or one with ATP permeability ^24, 25, 28^, an ATP release function of Panx1^1–89^ in the absence of wtPanx1 was tested. As shown in Figure 4, after an incubation period of 30 min, the ATP content of the supernatant of Panx1^1–89^ expressing cells was significantly higher than that of uninjected cells. This ATP release was boosted by a factor of ∼20, when the cells were incubated in high [K^+^] during the collection period. The Panx1 blocker carbenoxolone inhibited the ATP release.

**Figure 3.**
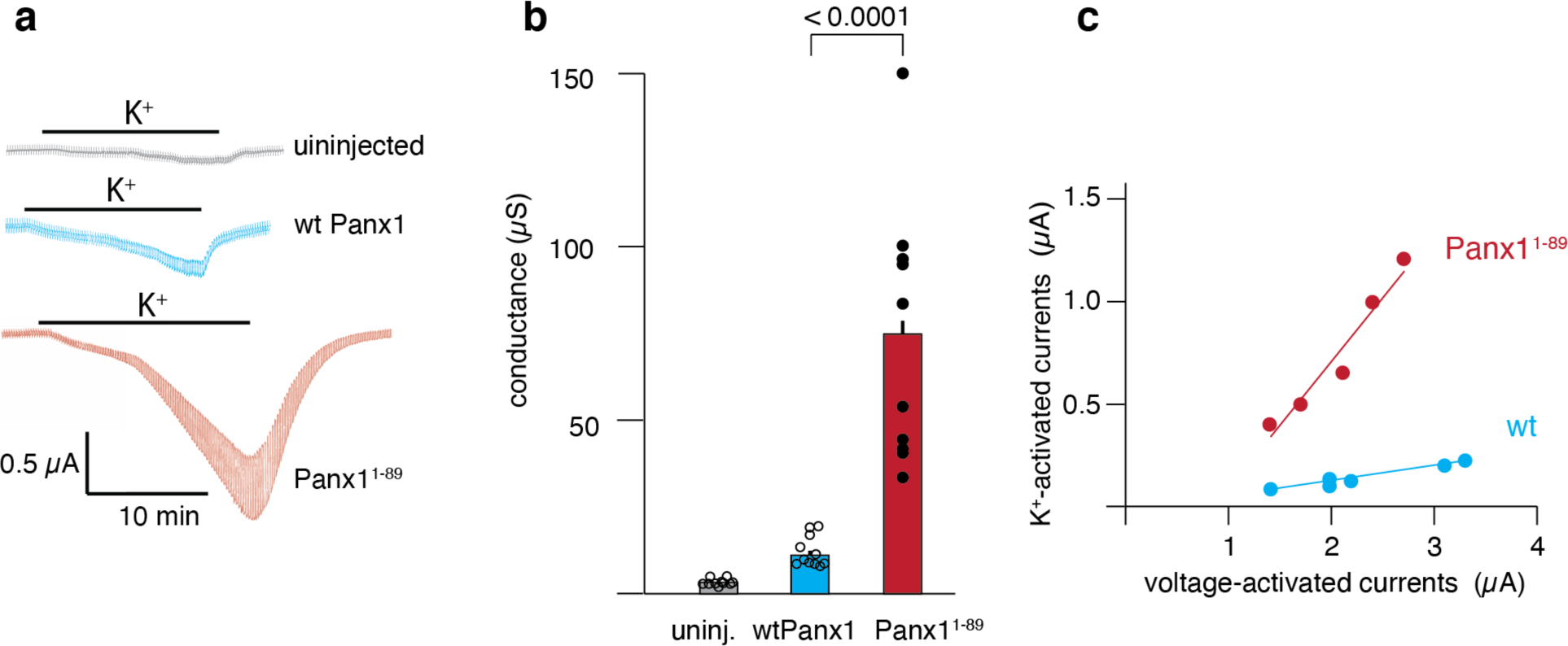
Effect of increased extracellular potassium ion concentration (85 mM, with K^+^ replacing Na^+^) on membrane currents in oocytes. **a**). Oocytes were voltage clamped at -60 mV and 12 mV depolarizing test pulses 5 seconds in duration were applied at 0.1 Hz. Traces for uninjected oocytes (top), oocytes expressing wtPanx1 (middle) or Panx1^1–89^ (bottom) are shown. K^+^ was applied as indicated by the horizontal bars. **b**) average calculated conductances for uninjected (grey), wtPanx1 (green) and Panx1^1–89^ (red) expressing oocytes are shown. Data are presented as mean ± SEM. n=10 (uninj.), 11 (wtPanx1) and 10 (P^1–89^). Only oocytes expressing wtPanx1 or Panx1^1–89^ beyond the threshold of 1 µA for voltage induced currents were analyzed. **c)** Relationship between K^+^-induced currents and currents induced by voltage steps from -60 to +60 mV as a measure of expression levels. The lines are based on regression analysis with the formula y=a+b*x yielding slopes of 0.62±0.09 (red) and 0.07±0.01 (green) and R-squares of 0.92 and 0.86, respectively.

**Figure 4.**
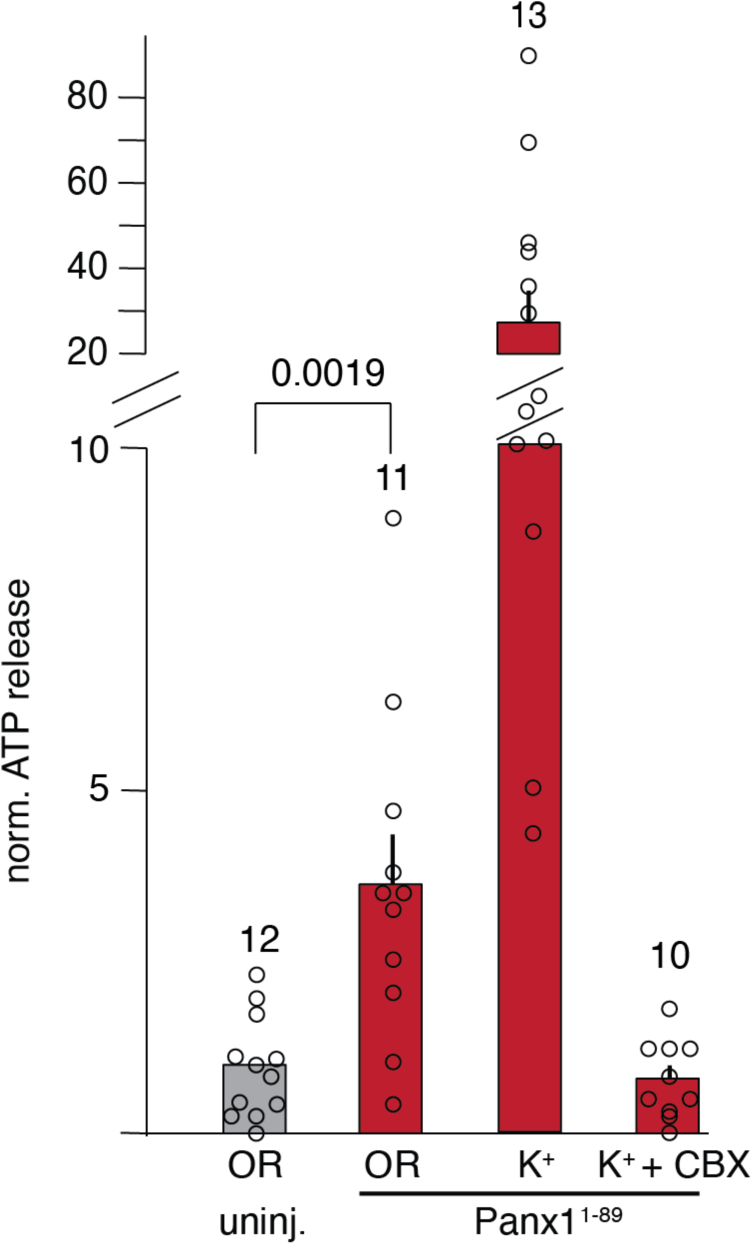
Release of ATP by uninjected oocytes (grey) and oocytes expressing Panx1^1–89^ (7 hours after injection of mRNA, red). Oocytes were incubated in Ringer solution (OR) or 85 mM K^+^ solution with and without CBX, as indicated. All data were normalized to the ATP content of the supernatant of unstimulated control oocytes. Data are presented as mean ± SEM. n is indicated above the bars. Data from 3 oocytes (2 from the Panx1^1–89^ OR and 1 from the K^+^+CBX data set) were excluded from analysis for visible damage to the cells.

Cells release ATP via exocytosis, via channels or in response to cell lysis. Since Panx1^1–89^ expressing cells have a limited life span, a contribution of lytic release of ATP cannot be ruled out. Therefore, as an independent method, we tested whether the Panx^1–89^ induced channel was able to pass other molecules in the size range of ATP such as cAMP.

To test cAMP permeability, we used the cystic fibrosis transmembrane conductance regulator (CFTR) channel, which is activated by high levels of intracellular cAMP. We reasoned that if the pore formed by Panx^1–89^ expression is large enough to pass cAMP, then it would provide a path for the influx of extracellular cAMP such that the levels of intracellular cAMP increase and in turn activate CFTR-mediated Cl^-^ flux out of cells co-expressing both Panx1^1–89^ and CFTR channels (Fig. 5a). Indeed, we found that application of extracellular cAMP activated a large inward current and a corresponding increase in membrane conductance in cells co-expressing CFTR and Panx1^1–89^ (Figure 5b). As a control, oocytes expressing either CFTR alone or Panx^1–89^ alone did not respond to 1 mM extracellular cAMP, despite a strong Forskolin-mediated inward current indicating robust membrane expression of CFTR (Fig. 5c, d). Figures 5 e and f show a quantitative analysis of the cAMP effects on currents and conductances. In summary, these data indicate that the large pore mediated by Panx1^1–89^ expression allows sufficient entry of extracellular cAMP into the cell to activate CFTR.

**Figure 5.**
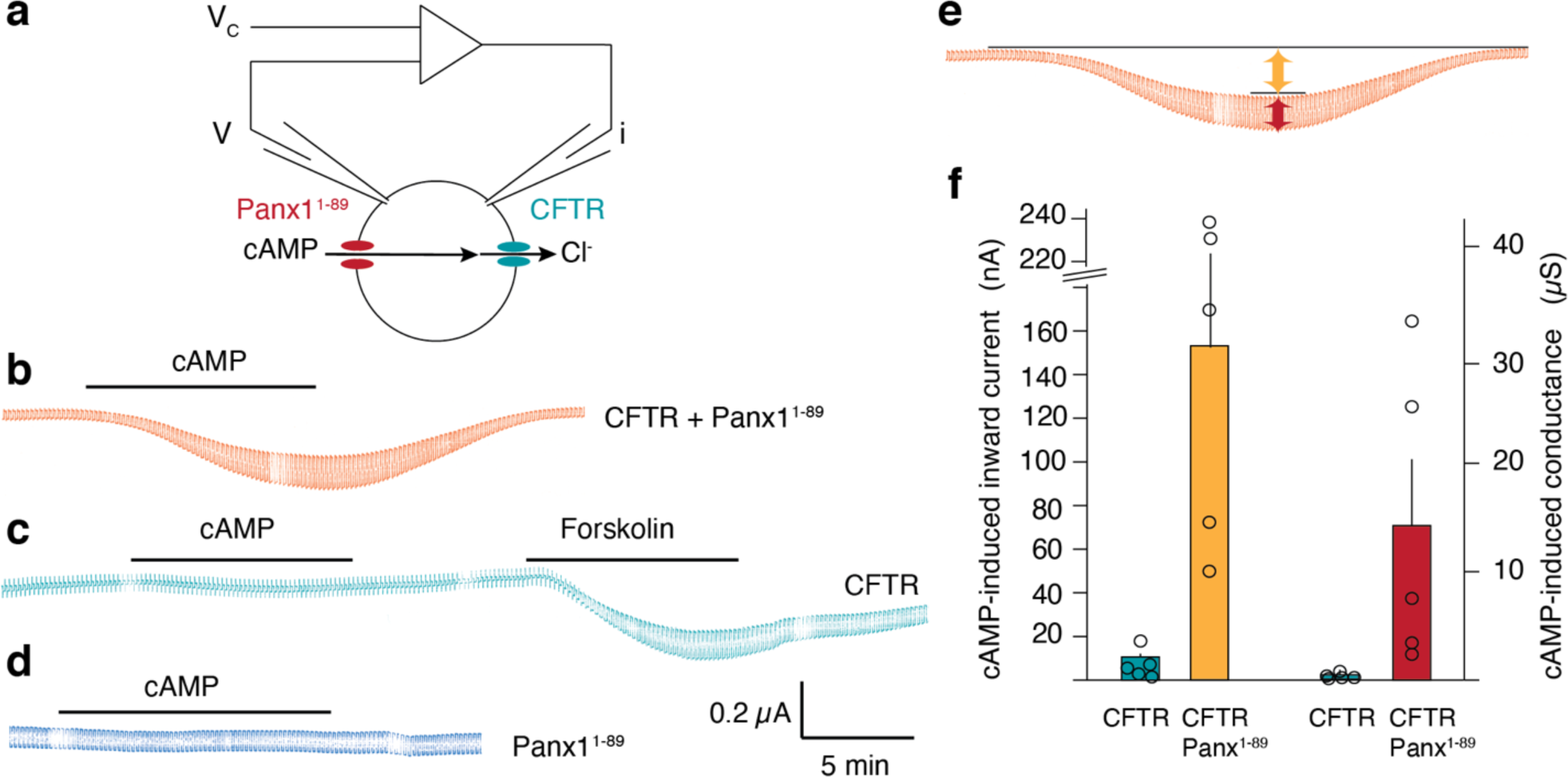
Uptake of extracellular cAMP. **a**) The schematic shows the voltage clamp arrangement and the deduced uptake of cAMP through Panx1^1–89^ mediated channels and subsequent activation of CFTR mediated chloride currents. Current traces are shown for oocytes co-expressing Panx1^1–89^ with CFTR (**b**), expressing CFTR alone (**c**) and expressing Panx1^1–89^ alone (**d**). 1 mM cAMP was applied extracellularly as indicated. To test for expression of CFTR, forskolin was applied as indicated in (**c**). Panel (**e**) shows how inward currents (orange arrow) and membrane conductance (red arrow) were determined. A quantitative analysis of cAMP-induced inward membrane currents and calculated conductances are shown in (**f**). The oocytes were voltage clamped at -60 mV and 12 mV depolarizing voltage steps were applied at a rate of 0.1 Hz. mRNAs for CFTR and for Panx1^1–89^ were injected 48 hours and 5 hours prior to the records, respectively. Data are presented as mean ± SEM. n=5.

### Panx1^1–89^ provides the pore lining of the induced channel

While the similarities in channel properties between wtPanx1 channels and the Panx1^1–89^ variant induced channels, i.e. inhibition of membrane currents by several Panx1 channel blockers, sensitivity to extracellular K^+^ and ATP permeability, suggest that Panx1^1–89^ by itself is sufficient to form the ion conducting pore of the channel, it cannot be excluded that the Panx1^1–89^ peptide associated with an endogenous protein and imposed Panx1 properties on it. To test whether Panx1^1–89^ provided the pore lining of the induced channel by itself, a cysteine substitution at position T62 (T62C) in Panx1^1–89^ was generated. It has previously been shown that the amino acid 62 is pore lining in the wtPanx1 channel ^42^, a conclusion confirmed by the cro-EM structure ^17–23^. Figure 6 shows the effect of the thiol reagent MTSET on currents in oocytes expressing the cysteine replacement mutant Panx1^1–89,T62C^. As observed for the equivalent mutation in wtPanx1, MTSET attenuated the currents carried by Panx1^1–89,T62C^ but not currents carried by Panx1^1–89^.

**Figure 6.**
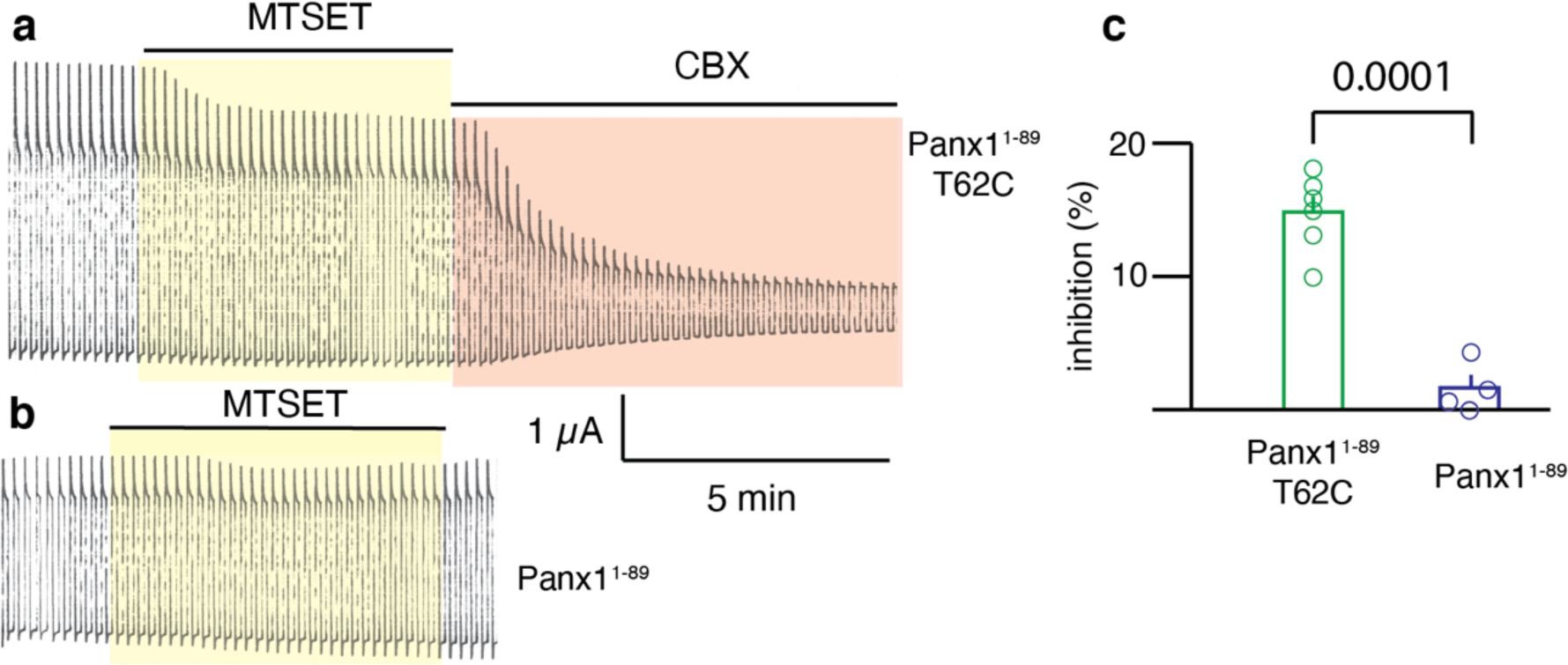
Effect of (2-(trimethylammonium)ethyl)MethaneThioSulfonate (MTSET) on membrane currents of the cysteine replacement mutant Panx1^1–89,T62C^. **a**) Oocytes were held at -60 mV and stepped to + 60 mV. 1mM MTSET was applied as indicated; this was followed by application of CBX to verify that the currents were mediated by Panx1^1–89^. **b**) As control, MTSET was applied to oocytes expressing Panx1^1–89^. **c**) Quantitative analysis of inhibition of membrane currents by MTSET in oocytes expressing Panx1^1–89,T62C^ (green) or Panx1^1–89^ (purple). Data are presented as mean ± SEM. n= 6 (green), 4 (purple).

Curiously, the effect of MTSET reversed after washout (Figure S4), as had also been observed for Panx1^T62C,C426S^ ^24^. In most other ion channels the effect of thiol reagents such as MTSET does not reverse unless reducing agents are employed. However, it has been shown for large pore channels, such as channels formed by connexins, the effect of thiol reagents reverses spontaneously upon withdrawal of the thiol reagent ^24, 43–45^. This reversibility has been explained by the ability of cytoplasmic reducing agents, in particular glutathione, to enter the large pore and getting access to the modified cysteine in the channel pore. Thus, the reversibility of the MTSET effect on Panx1^1–89,T62C^ is a further indication that the channel formed by this peptide is in a large pore conformation similar to the one observed in wtPanx1 in response to select stimuli.

Wild type Panx1 contains four conserved extracellular cysteines, two in each of the two extracellular loops. Mutation of any of these cysteines leads to loss of channel function, suggesting a key structural/functional role of these moieties ^46^. In the Panx1^1–89^ peptide two of these four cysteines are retained. To test whether these cysteines are disulfide bonded, the reducing agent TCEP was applied. As shown in Figure 7a and b, 10 mM of TCEP attenuated the Panx1^1–89^ currents by 45.7 ± 5.6 %. Subsequent addition of the thiol reagents MPB or MTSET (Figure 7b) did not further affect the currents, indicating that the cysteines do not contribute to the pore lining. The spontaneous ATP release by Panx1^1–89^ expressing oocytes was nearly abolished by the addition of TCEP (Figure 7c). The large effect on ATP release with a more than 50% residual conductance after TCEP application suggests a change in permeability, so that the flux of larger molecules is more affected than that of smaller ions (Figure S5). The published cryo-EM structures of wild type hPANX1 show the cysteines disulfide bonded between the two extracellular loops (C66 to C265 and C84 to C246) ^21, 22^. Such bonding is not possible in Panx1^1–89^. However, the two cysteines in the first extracellular loop are in close proximity. Thus, in the Panx1^1–89^, peptide the two cysteines appear to be well positioned for disulfide bonding.

**Figure 7.**
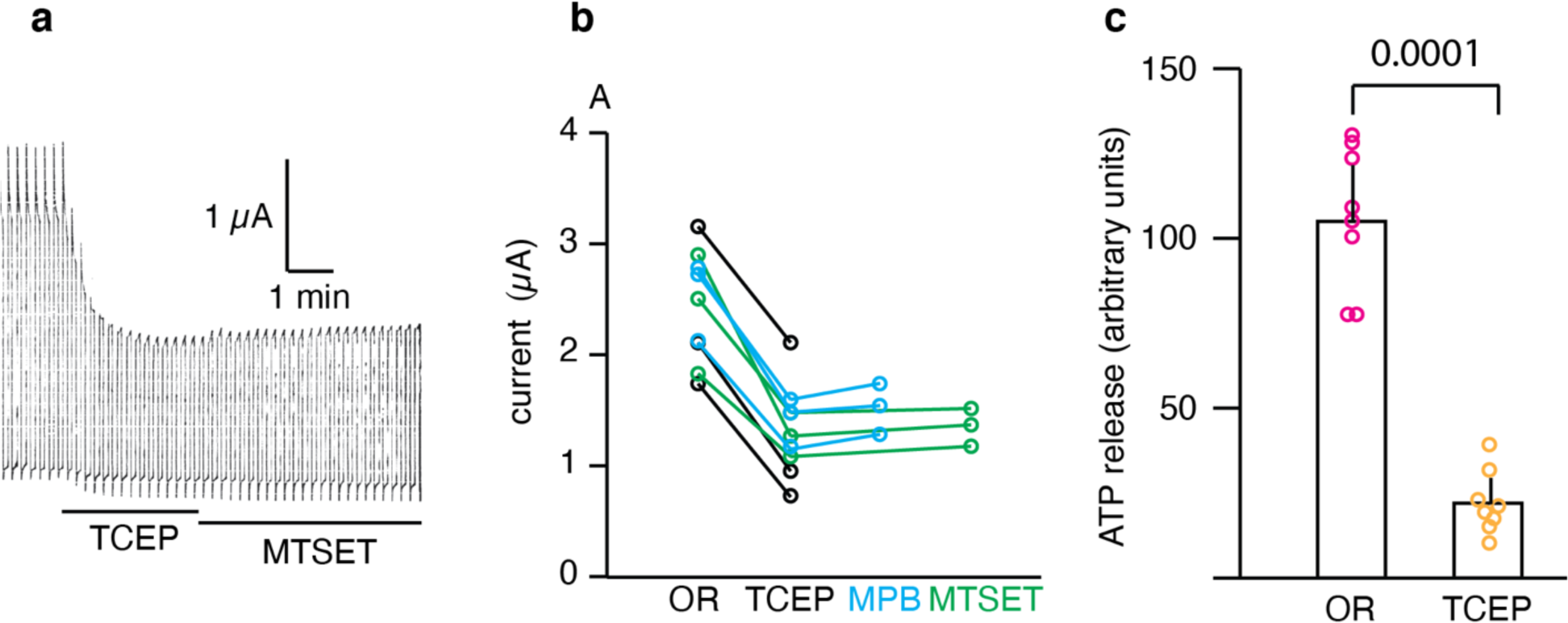
Effect of the reducing agent TCEP on Panx1^1–89^-mediated membrane currents and ATP release. **a)** Traces of membrane currents induced by voltage steps from -60 to +60 mV in an oocyte expressing Panx1^1–89^. TCEP was applied as indicated and was followed by the thiol reagent MTSET. **b)** Membrane currents recorded from 9 oocytes expressing Panx1^1–89^ before and after application of TCEP black, blue and green symbols and lines. In 3 oocytes TCEP was followed by the thiol reagents MPB (blue) or MTSET (green). **c)** Spontaneous ATP release of oocytes expressing Panx1^1–89^ incubated in oocyte Ringer solution (OR) or in OR supplemented with TCEP. Since TCEP attenuates the luciferase reaction used for ATP determination, the supernatant from the oocytes in OR were spiked with an equimolar amount of TCEP. Uninjected oocytes served as control and the mean values were subtracted from the values of the Panx1^1–89^ expressing oocytes. Data are presented as mean ± SEM, n=8.

The reducing effect of TCEP not only attenuated Panx1^1–89^ currents and ATP release, but also abolished the response of Panx1^1–89^ to increased extracellular [K^+^]. Figure 8 shows that in the presence of TCEP, application of 80 mM [K^+^] did not increase the holding current or the membrane conductance. Changes in both parameters were typically observed in the non-reduced Panx1^1–89^ channel (compare to Figure 3). These data indicate that reduction of the disulfide bonds in Panx1^1–89^ results in a major conformational change of the extracellular channel entry (Figure S5), which affects conductance, permeability and activation by K^+^.

**Figure 8.**
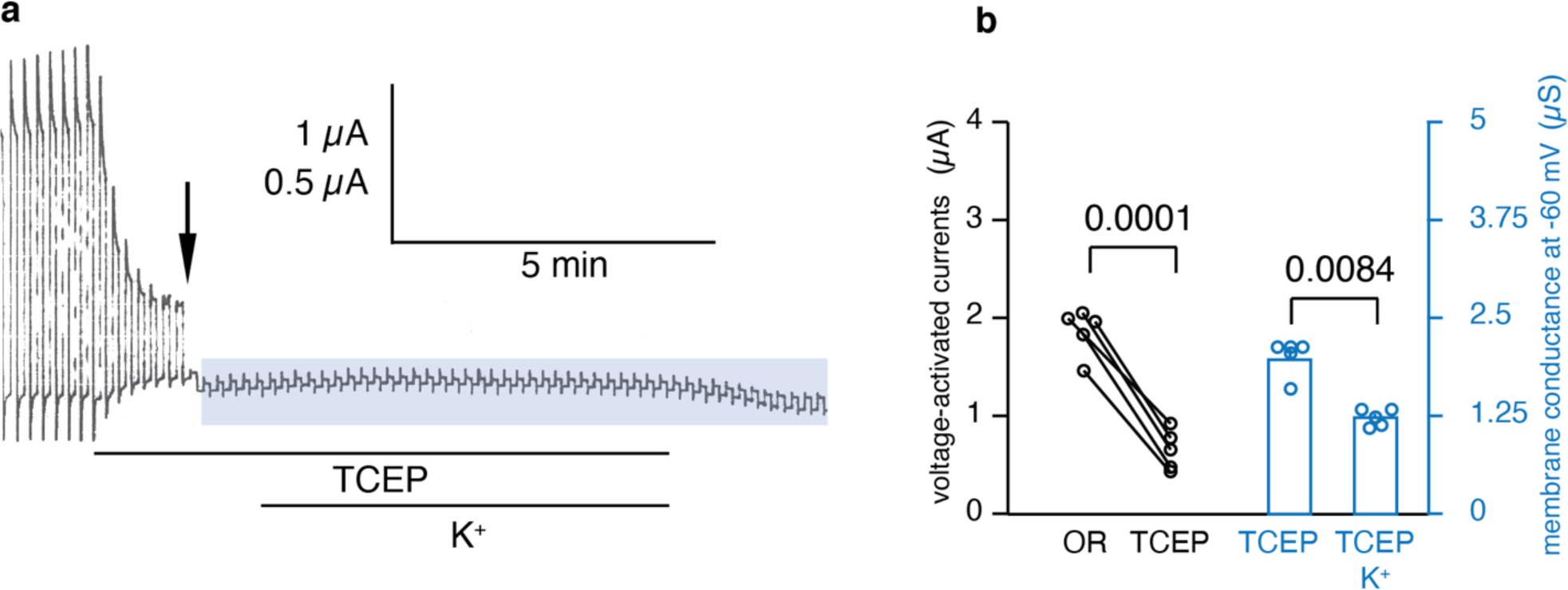
Loss of activation of Panx1^1–89^ by extracellular K^+^ in response to reduction by TCEP. **a)** Current traces of an oocyte expressing Panx1^1–89^. The membrane potential was clamped at – 60 mV and voltage steps to +60 mV were applied at a rate of 0.1 Hz. 10 mM TCEP was applied as indicated by the bar. At the time point indicated by the arrow, the pulse protocol was changed to 12 mV pulses from a -60 mV holding potential to test the effect of K^+^ on membrane conductance. 85 mM K^+^ was applied in the presence of TCEP as indicated by the bar. Scale: 1 µA for 120 mV pulses and 0.5 µA for 12 mV pulses. **b)** Quantitative analysis of the effect of TCEP on 120 mV voltage step induced membrane currents (left) and membrane conductance during the 12 mV voltage step period (right). In contrast to non-reducing conditions (compare Figure 3), K^+^ did not activate Panx1^1–89^. The attenuation in membrane conductance during the application of K^+^, could be due to a continued reduction of Panx1^1–89^ activity by TCEP or a negative effect of K^+^ on membrane conductance under reducing conditions. Data for conductance are presented as mean ± SEM, n=5.

## DISCUSSION

Based on the correlation between the expression of a severely truncated Panx1 mutant, Panx1^1–89^, with the metastatic potential of breast cancer cells and the well established role of extracellular ATP in metastasis of several types of cancers, *Furlow et al.* ^14^ formulated an intriguing hypothesis for a Panx1 role in metastatic cell survival in the microvasculature. It has been suggested that the Panx1^1–89^ peptide, amplifies the release of ATP by the membrane channel formed by wtPanx1. Since measurements of ATP release were the sole basis for this conclusion, we set out to perform a more detailed characterization of biophysical properties of these presumably heteromeric channels in the *Xenopus* oocyte expression system. However, control oocytes expressing exclusively Panx1^1–89^ exhibited a membrane conductance as soon as three hours after injection of the mRNA. This conductance was inhibited by a series of inhibitors of the wtPanx1 channel, including probenecid, carbenoxolone, BB FCF and BzATP. Thus, it appears that Panx1^1–89^ may not require the presence of wtPanx1 to render a membrane conductance. To do this, the Panx1^1–89^ peptide could have associated with an endogenous membrane channel or it may have formed a channel by itself, probably by entering an oligomeric state.

It is unlikely, if not impossible, that a peptide with a single transmembrane segment forms a membrane pore in a monomeric form. Instead, viroporins with a single transmembrane segment form pores by oligomerizing 4,5 or 6 subunits ^30–33^. The wtPanx1 channel is known to assemble as a heptamer ^17–23^. Presently, it is not known whether the channel formed by Panx1^1–89^ is a tetramer, a pentamer, a hexamer, a heptamer like wtPanx1 or even assumes a higher oligomeric state. It is also possible that Panx1^1–89^ assembles into various oligomeric states, since elements controlling the oligomerization of the wt channel possibly are missing in the peptide. Thus, different oligomeric states may co-exist.

When co-expressed with wtPanx1, as in cancer cells, it has to be assumed that in the absence of specific controlling factors a spectrum of heteromeric and homomeric assemblies are to be encountered. In metastatic cancer cells the ratio of wt over mutant mRNA was about 3:1. The observation that in oocytes mutant channels appeared much earlier than wt channels, on the other hand, suggests a higher translation rate of the shorter peptide. Consequently, the distribution of the various heteromers and homomers cannot be assessed at this time. Nevertheless, it can be safely assumed that a certain (albeit small) percentage of co-expressing cells will contain homomeric Panx1^1–89^ channels.

Although the truncation deletes 80 % of the Panx1 protein, it is not totally surprising that Panx1^1–89^ could form a functional membrane channel. The 1-89 sequence contains most, if not all, of the pore-lining amino acids of the wt channel ^17–22, 42^. In addition, the “binding sites” for channel regulators or part of them located in the first extracellular loop of the wt channel are still contained in Panx1^1–89^ ^17, 35–38, 47^ . While it is conceivable that Panx1^1–89^ may associate with a channel endogenous to oocytes, the finding that the conductance mediated by Panx1^1–89^ is inhibited by thiol reagents in a cysteine mutant, Panx1^1–89, T62C^, similar to the inhibition of Panx1^T62C,C426S^ by the same thiol reagent, suggests that Panx1^1–89^ by itself forms the channel pore. There is precedence for single membrane spanning peptides to form functional membrane channels, as shown for viroporins or alamethicin ^30–33, 48, 49^.

Since oocytes did not survive longer than 12 hours after injection of mRNA for Panx1^1–89^, there may be a simple explanation for the failure to observe ATP release by Panx1^1–89^ alone expressed in HEK cells in the initial paper on the truncation mutant found in metastatic cells. HEK cells cells that did express Panx1^1–89^ at high levels might not have survived and the HEK cells surviving transfection may not have expressed Panx1^1–89^ or at reduced rate. The observation that oocytes co-injected with Panx1^1–89^ and wtPanx1 mRNA survived several days suggests that wtPanx1 is protective against the effects of Panx1^1–89^ and that Panx1^1–89^ and wtPanx1 proteins interact with each other. For example, the channels formed by these proteins could be heteromeric and exhibit a lower open probability and /or a smaller pore size.

Alternatively, the Panx1^1–89^ peptide may not shuttle to the cell membrane in HEK cells. For example, proper folding may be temperature sensitive. Consequently, the peptide may fold properly in *Xenopus* oocytes at room temperature but not in HEK cells at 37° C. wtPanx1 in that scenario may act as a chaperone and allow Panx1^1–89^ to shuttle properly folded to the plasma membrane. A prime example for such a scenario is the CFTR ΔF508 mutation, which expresses functionally in *Xenopus* oocytes but is retained in the ER in tissue culture cells at 37° C because of misfolding ^50–52^.

It is well documented that truncation of the carboxyterminal 47 amino acids by caspase cleavage of Panx1 renders the channel constitutively active ^39^. In this respect the truncation by caspase by 48 (mouse) or 47 (human) amino acids and the severe truncation of Panx1^1–89^, which deletes 337 amino acids have similar consequences. However, while the caspase-cleaved wtPanx1 channel in the absence of an additional stimulus is a highly selective Cl^-^ channel without ATP permeability ^16, 25, 40^, the channel formed by Panx1^1–89^ was found to exhibit a different permeability as indicated by the shift of the reversal potential versus wtPanx1, the release of ATP and the uptake of cAMP without an additional stimulus. Thus, the Panx1^1–89^ in the absence of a stimulus is not only constitutively active but also adopts a conformation akin to the large pore formed by wtPanx1 in response to a variety of physiological or pathological stimuli.

Another similarity between wtPanx1 and Panx1^1–89^ channels is their response to increased extracellular potassium ion concentration. Several amino acids involved in the activation of wtPanx1 are still contained in the truncated Panx1^1–89^. However, the response of Panx1^1–89^ was considerably stronger than that of wtPanx1, indicating that constraints to the K^+^ activation are absent in the truncated protein. A boost to the K^+^ response has been previously shown for two alanine replacement mutants of the intact Panx1 protein (D241A and L266A), consistent with such a loss of constraints. Although the concentration of K^+^ required to activate wtPanx1 is well beyond the physiological range, it is conceivable that under pathological conditions, such as the penumbra of a stroke lesion ^53^, sufficiently high [K^+^] is reached. In the case of Panx1^1–89^, even slight elevations of [K^+^] may result in large amounts of ATP being released.

The present findings do not rule out that in metastatic cells, wtPanx1 and Panx1^1–89^ form heteromeric channels that are constitutively active and permeant to ATP as proposed ^14^. Instead, the present results show that Panx1^1–89^ can form functional membrane channels and mediate ATP release even in the absence of wtPanx1. Most importantly, the homomeric Panx1^1–89^ are susceptible to inhibition by the same drugs that inhibit wtPanx1 channels. Thus, compounds such as probenecid should be considered for complementary use to standard treatments of metastatic cancers if the Panx1^1–89^ mutants are present.

Another aspect of the present findings is that the Panx1^1–89^ mutant may simplify the structure-function analysis of Panx1 channels. Since most basic aspects of the channel are conserved in the heavily truncated peptide, the pore structure analysis should be facilitated. Furthermore, the observation that the Panx1^1–89^ channel appears to be constitutively active in the so far elusive “large pore” conformation may enhance the chance to capture this conformation by cryo-EM.

## MATERIAL AND METHODS

### Materials

Mouse pannexin1 was kindly provided by Dr. Rolf Dermietzel (University of Bochum). Human CFTR was kindly provided by Dr. Seth Alper (Harvard Medical School).

ATP, BzATP, Brilliant Blue FCF (BB FCF), cAMP, carbenoxolone, forskoline and tris(2- carboxyethyl)phosphine (TCEP) were purchased from Sigma Aldrich. Probenecid was obtained from Alfa Aesar. The thiol reagents MTSET and MPB were purchased from Toronto Research Chemicals and Sigma Aldrich, respectively.

### Mutagenesis

The Panx1 mutants were engineered with QuickChange II site-directed mutagenesis kit (Stratagene) according to the manufacturer’s specifications. The purified mutant plasmids were sequenced by Genewiz.

The following primers were used:

Stop codon following Q89:

5’-GCTGCTGTACAGTAGAAGAGCTCCCTGC-3’

5’-GCAGGGAGCTCTTCTACTGTACAGCAGC-3’

Cysteine replacement T62C:

5’-GGAGATCTCCATCGGTTGTCAGATAAGCTGC-3’

5’-GCAGCTTATCTGACAACCGATGGAGATCTCC-3’

### Preparation of oocytes

All procedures were approved by the University of Miami Institutional Animal Care and Use Committee and conducted in accordance with the Guiding Principles for Research Involving Animals and Human Beings of the American Physiological Society. Ovaries were harvested from adult female *Xenopus laevis*. Ovaries were cut into small pieces and incubated in collagenase (2.5 mg/ml; Worthington) in calcium-free oocyte Ringer (OR) solution, stirring at one turn per second at room temperature. Typically, the incubation period was 3 h for oocytes to be separated from the follicle cells. After thorough washing with regular OR (82.5 mM NaCl, 2.5 mM KCl, 1 mM Na_2_HPO_4_, 1 mM MgCl_2_, 1 mM CaCl_2_, and 5 mM HEPES), oocytes devoid of follicle cells and having a uniform pigmentation were selected and stored in OR at 18°C for 18 h to 3 days before electrophysiological analysis at room temperature.

### Preparation of mRNA and electrophysiology

The plasmid containing mouse pannxin1 or its mutants in pCS2 was linearized with Not I. The CFTR cDNA was linearized with XhoI, In vitro transcription was performed with SP6 (Panx1) or T7 (CFTR) polymerase, using the Message Machine kit (Ambion, Austin, TX). mRNAs were quantified by absorbance (260 nm), and the proportion of full-length transcripts was checked by agarose gel electrophoresis. In vitro-transcribed mRNAs (60 nl for wt and 32 nl for Panx1^1–89^ at ∼1µg/µl) were injected into *Xenopus* oocytes. Cells were kept in regular Ringer solution with the antibiotic streptomycin (10 mg/ml). Whole cell membrane currents of oocytes were measured using a two-electrode voltage clamp (Gene Clamp 500B, Axon Instruments/Molecular Devices) under constant perfusion according to the protocols described in the figures. Glass pipettes were pulled using a P-97 Flaming/Brown puller (Sutter). Two electrophysiological protocols were used: to determine membrane conductance, the membrane potential was held at -60 mV and small test pulses lasting 6 s to -48 mV were applied at a rate of 0.1 Hz; alternatively, voltage ramps lasting 35 s were applied, typically from -100 mV to +100 mV.

Because of the abbreviated life time of oocytes expressing the mutant Panx1^1–89^, only a short time window of 4 to 8 hours after mRNA injection was available for electrical recording and determination of ATP release. In contrast, adequate expression of of CFTR required ∼48 hours after injection of mRNA. For co-expression of CFTR and Panx1^1–89^, CFTR mRNA was injected first, followed by injection of Panx1^1–89^ 48 hours later. After 5-7 hours, co-expressing cells were subjected to voltage clamp analysis.

### ATP release assay

ATP flux was determined by luminometry. Oocytes, 4 hours after injection of Panx1^1–89^ mRNA or two days after injection of wtPannexin1 were analyzed. Ninety microliters of the oocyte supernatant were added to 40 µl of luciferase-luciferin solution (Promega, Madison, WI) for assaying luciferase activity. Data from oocytes with visible damage were excluded from analysis.

## SUPPLEMENTARY MATERIALS

Supplementary Material contains 5 Supplementary Figures.

**Supplementary Figure 1**. Photographs of *Xenopus* oocytes 3 days after harvesting from the ovaries.

**Supplementary Figure 2**. Membrane topology of wtPanx1 and Panx1^1–89^ channels.

**Supplementary Figure 3**. Current traces of oocytes injected with mRNA for Panx1^1–89^ 3 to 5 hours prior to the recording.

**Supplementary Figure 4**. Reversibility of the effect of MTSET on membrane channels in oocytes expressing Panx1^1– 89,T62C^.

**Supplementary Figure 5**. Schematic illustrating the effect of the reducing agent TCEP on the pore conformation of Panx1^1–89^.

## ACKNOWLEDGEMENTS

We thank Drs. H. Peter Larsson and Kenneth Muller for critically reading the manuscript.

## AUTHOR CONTRIBUTION

GD and RB-S conceived the project and supervised all research. CM, GD, and RB-S wrote the manuscript. CM, GD, and RB-S designed the experiments. JW performed the experiments. JW, GD and RB- S analyzed data. JW, GD and RB-S designed and performed statistical analysis.

---

**S1.**
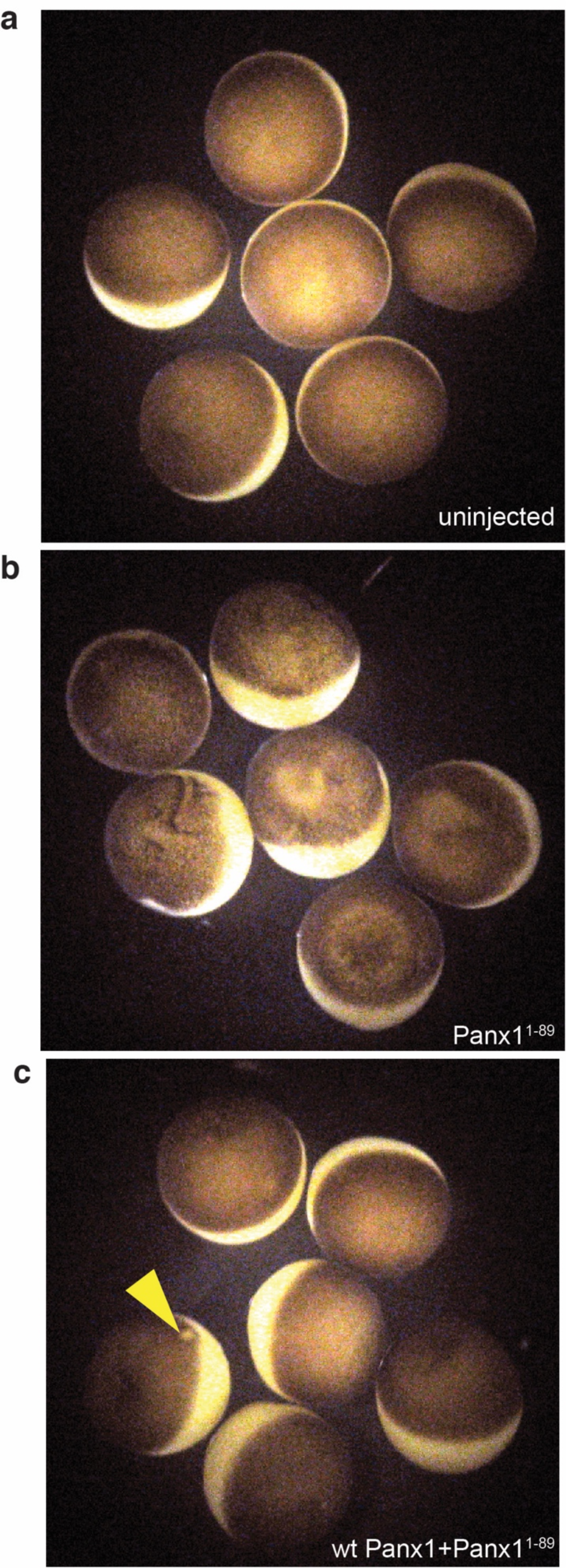
Photographs of *Xenopus* oocytes 3 days after harvesting from the ovaries. **a**) uninjected control oocytes exhibited even pigmentation at the animal pole and were well rounded. **b**) Oocytes injected 12 hours prior with mRNA encoding Panx1^1–89^ exhibited uneven pigmentation, indentations of the membrane and signs of shrinkage. **c**) Oocytes co-injected with wtPanx1 and Panx1^1–89^ mRNA (75 ng each in a volume of 60 nl) 24 hours prior were indistinguishable from control oocytes, except one cell showing an injection scar (arrowhead).

**S2.**
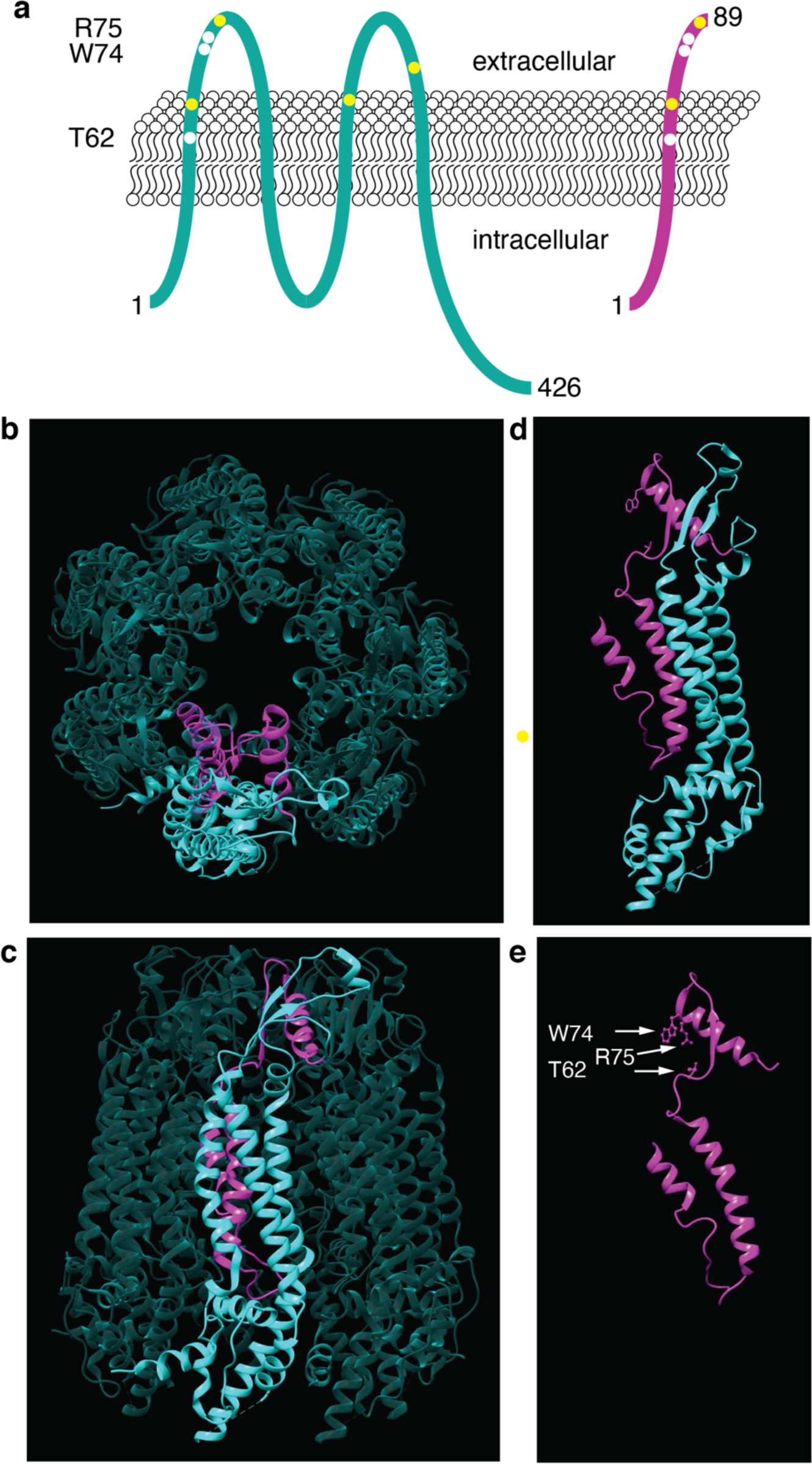
Membrane topology of wtPanx1 and Panx1^1–89^ channels. (a) Schematic of membrane topology of wtPanx1 (green) and Panx1^1–89^ (purple) with amino acids engaged in pore lining (including external constriction i.e. W74 and R75) and binding of channel modulators marked by white dots. Yellow dots indicate the two conserved cysteines in the extracellular loops. Structure of the peptide within the wtPanx1 model based on the cryo-EM structure (pdb ID 7F8J) ^54^. For better visibility, one protomer is highlighted. The color scheme from (a) is used to identify Panx1^1–89^ (b) top view and (c) side view of wtPanx1. (d) side view of the protomer and (e) side view of the amino acids within the protomer with important (W74, R75 and T62) residues as stick-and-ball representation. The figure was prepared Chimera. ^55^

**S3.**
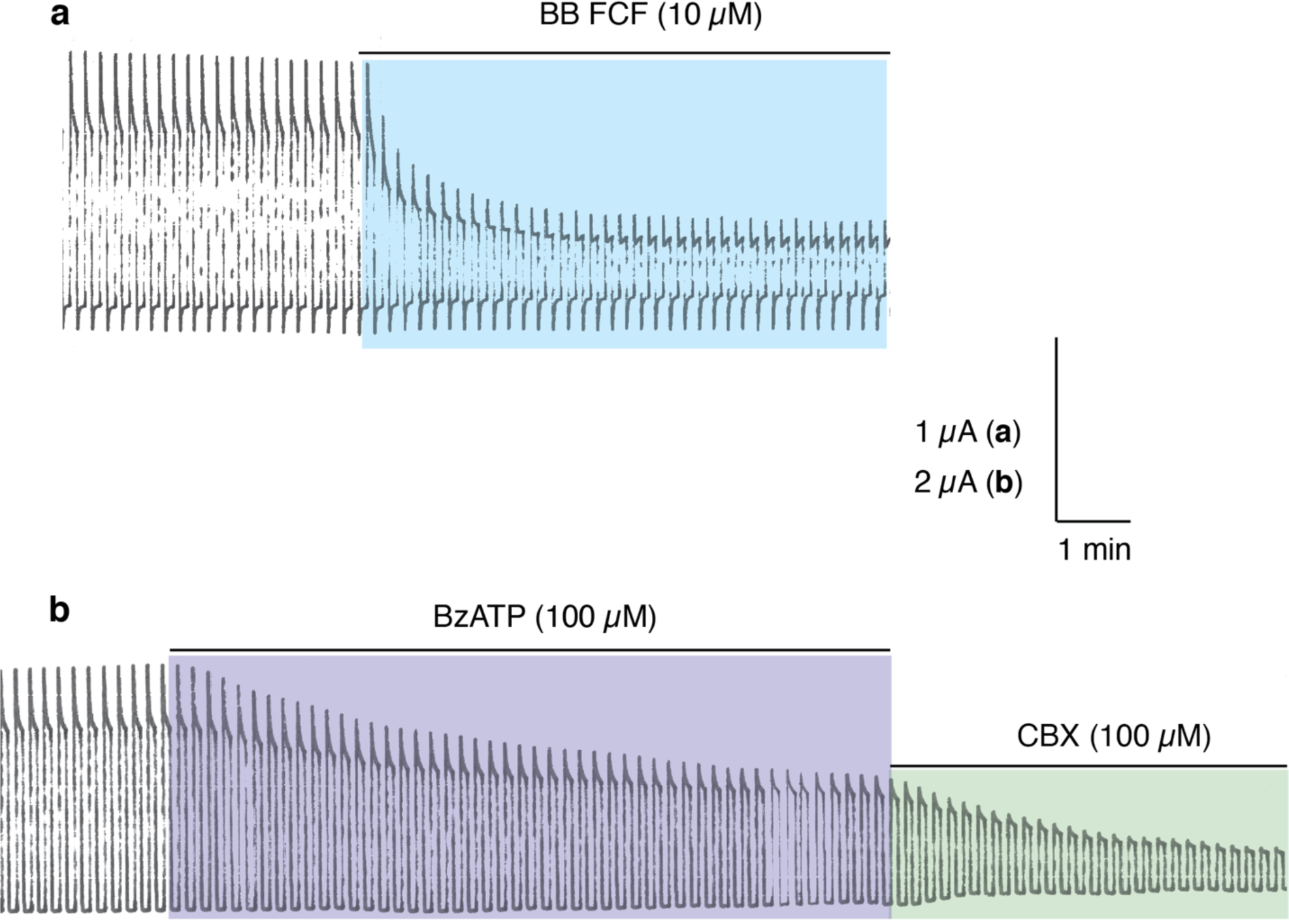
Current traces of oocytes injected with mRNA for Panx1^1–89^ 3 to 5 hours prior to the recording. **a**) The food dye BB FCF applied at 10 µM attenuated the currents induced by voltage steps from -60 to +60 mV. **b**) BzATP at 100 µM also partially attenuated the currents, which were further inhibited by 100 µM carbenoxolone (CBX).

**S4.**
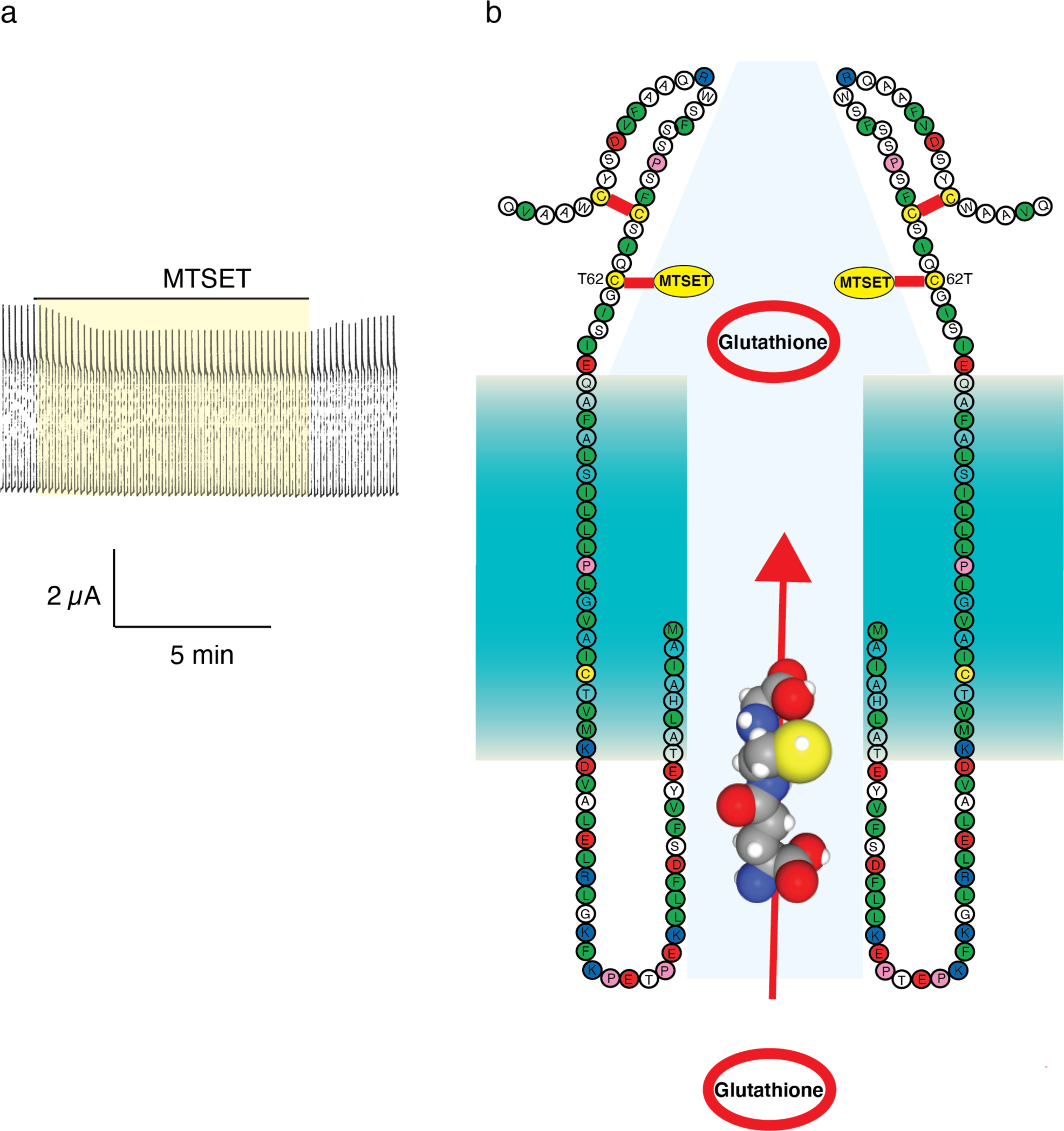
Reversibility of the effect of MTSET on membrane channels in oocytes expressing Panx1^1–89,T62C^. Since the reversal of MTSET in other ion selective channels with substituted cysteines typically requires reducing agents, this observation suggests that endogenous cytoplasmic reducing agents such as glutathione reached the 62 position in the channel induced by Panx1^1-89,T62C^.

**S5.**
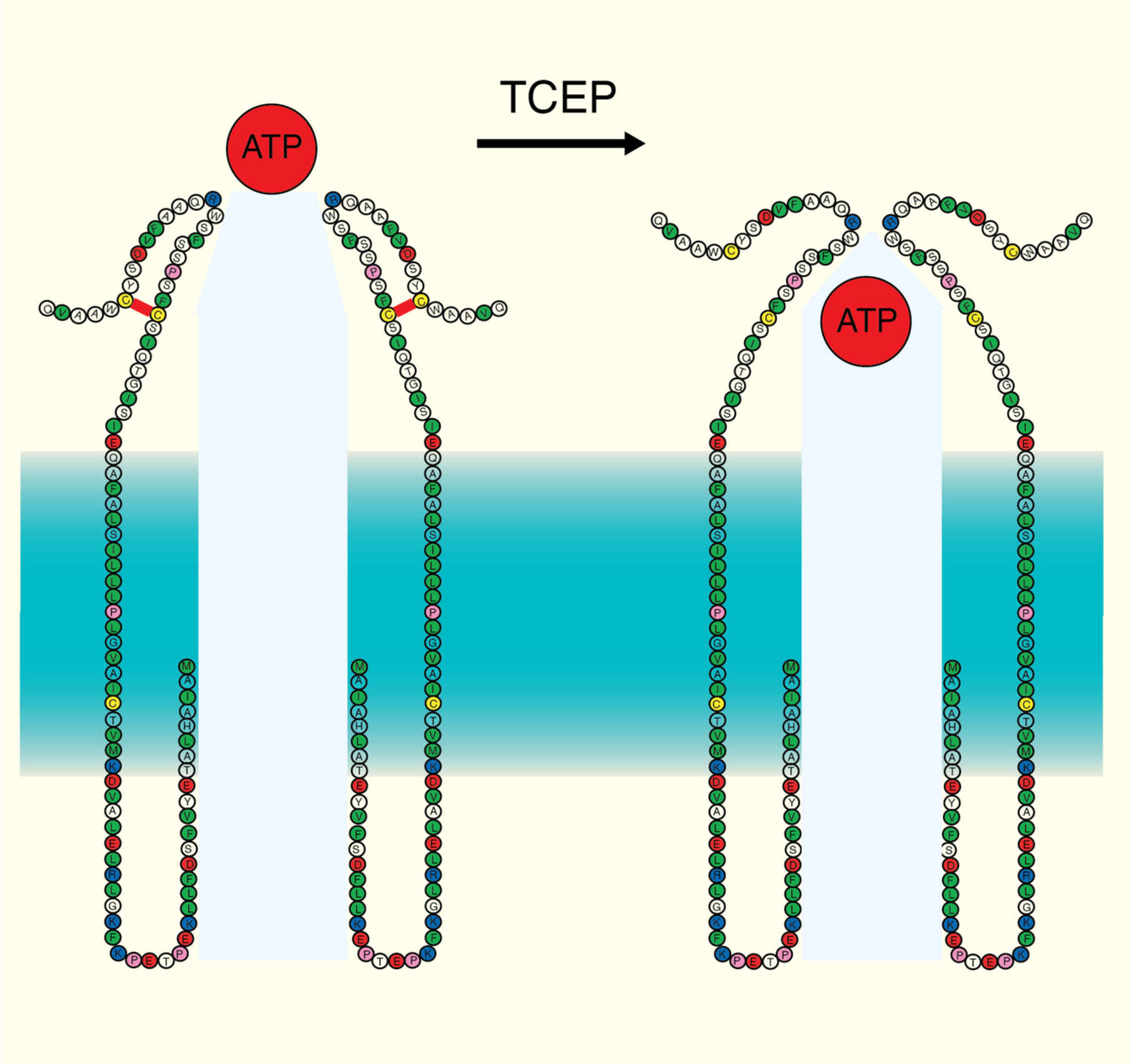
Schematic illustrating the effect of the reducing agent TCEP on the pore conformation of Panx1^1–89^. All presently available Panx1 cryo-EM structures show W74 and R75 as a constriction of the external aspect of the channel pore thereby governing permeability of the channel. In Panx1^1–89^ two of the four conserved extracellular cysteines are retained. These cysteines are likely to be disulfide bonded as the reducing agent TCEP significantly reduced membrane currents and ATP release. It appears that ATP release was more affected than membrane currents by TCEP, suggesting a change in permeability. This change in conformation also rendered the channel insensitive to extracellular K^+^.

## REFERENCES

1. Vultaggio-Poma, V., Sarti, A.C. & Di Virgilio, F. Extracellular ATP: A Feasible Target for Cancer Therapy. Cells 9 (2020).

2. Brenet, M. et al. Thy-1 (CD90)-Induced Metastatic Cancer Cell Migration and Invasion Are beta3 Integrin-Dependent and Involve a Ca(2+)/P2X7 Receptor Signaling Axis. Front Cell Dev Biol 8, 592442 (2020).

3. Martin, S. et al. An autophagy-driven pathway of ATP secretion supports the aggressive phenotype of BRAF(V600E) inhibitor-resistant metastatic melanoma cells. Autophagy 13, 1512–1527 (2017).

4. Kepp, O. et al. ATP and cancer immunosurveillance. EMBO J 40, e108130 (2021).

5. Ledderose, C. et al. Cutting off the power: inhibition of leukemia cell growth by pausing basal ATP release and P2X receptor signaling? Purinergic Signal 12, 439–451 (2016).

6. Penuela, S. et al. Loss of pannexin 1 attenuates melanoma progression by reversion to a melanocytic phenotype. J Biol Chem 287, 29184–29193 (2012).

7. Freeman, T.J. et al. Inhibition of Pannexin 1 Reduces the Tumorigenic Properties of Human Melanoma Cells. Cancers (Basel) 11 (2019).

8. Sayedyahossein, S. et al. Pannexin 1 binds beta-catenin to modulate melanoma cell growth and metabolism. J Biol Chem 296, 100478 (2021).

9. Bao, L., Sun, K. & Zhang, X. PANX1 is a potential prognostic biomarker associated with immune infiltration in pancreatic adenocarcinoma: A pan-cancer analysis. Channels (Austin) 15, 680–696 (2021).

10. Zuzul, M. et al. The Expression of Connexin 37, 40, 43, 45 and Pannexin 1 in the Early Human Retina and Choroid Development and Tumorigenesis. Int J Mol Sci 23 (2022).

11. Liu, H. et al. In vitro effect of Pannexin 1 channel on the invasion and migration of I-10 testicular cancer cells via ERK1/2 signaling pathway. Biomed Pharmacother 117, 109090 (2019).

12. Shi, G. et al. Panx1 promotes invasion-metastasis cascade in hepatocellular carcinoma. J Cancer 10, 5681–5688 (2019).

13. Penuela, S., Gehi, R. & Laird, D.W. The biochemistry and function of pannexin channels. Biochim Biophys Acta 1828, 15–22 (2013).

14. Furlow, P.W. et al. Mechanosensitive pannexin-1 channels mediate microvascular metastatic cell survival. Nat Cell Biol 17, 943–952 (2015).

15. Syrjanen, J., Michalski, K., Kawate, T. & Furukawa, H. On the molecular nature of large-pore channels. J Mol Biol 433, 166994 (2021).

16. Mim, C., Perkins, G. & Dahl, G. Structure versus function: Are new conformations of pannexin 1 yet to be resolved? J Gen Physiol 153 (2021).

17. Michalski, K. et al. The Cryo-EM structure of pannexin 1 reveals unique motifs for ion selection and inhibition. Elife 9 (2020).

18. Deng, Z. et al. Cryo-EM structures of the ATP release channel pannexin 1. Nat Struct Mol Biol 27, 373–381 (2020).

19. Jin, Q. et al. Cryo-EM structures of human pannexin 1 channel. Cell Res 30, 449–451 (2020).

20. Mou, L. et al. Structural basis for gating mechanism of Pannexin 1 channel. Cell Res 30, 452–454 (2020).

21. Qu, R. et al. Cryo-EM structure of human heptameric Pannexin 1 channel. Cell Res 30, 446–448 (2020).

22. Ruan, Z., Orozco, I.J., Du, J. & Lu, W. Structures of human pannexin 1 reveal ion pathways and mechanism of gating. Nature (2020).

23. Zhang, S. et al. Structure of the full-length human Pannexin1 channel and insights into its role in pyroptosis. Cell Discov 7, 30 (2021).

24. Wang, J. et al. The membrane protein Pannexin1 forms two open-channel conformations depending on the mode of activation. Sci Signal 7, ra69 (2014).

25. Wang, J. & Dahl, G. Pannexin1: a multifunction and multiconductance and/or permeability membrane channel. Am J Physiol Cell Physiol 315, C290–C299 (2018).

26. Romanov, R.A. et al. The ATP permeability of pannexin 1 channels in a heterologous system and in mammalian taste cells is dispensable. J Cell Sci 125, 5514–5523 (2012).

27. Ma, W. et al. Pannexin 1 forms an anion-selective channel. Pflugers Arch 463, 585–592 (2012).

28. Dahl, G. ATP release through pannexon channels. Philos Trans R Soc Lond B Biol Sci 370 (2015).

29. Dahl, G. The Pannexin1 membrane channel: distinct conformations and functions. FEBS Lett 592, 3201–3209 (2018).

30. Nieva, J.L., Madan, V. & Carrasco, L. Viroporins: structure and biological functions. Nat Rev Microbiol 10, 563–574 (2012).

31. Surya, W., Li, Y., Verdia-Baguena, C., Aguilella, V.M. & Torres, J. MERS coronavirus envelope protein has a single transmembrane domain that forms pentameric ion channels. Virus Res 201, 61–66 (2015).

32. Largo, E., Queralt-Martin, M., Carravilla, P., Nieva, J.L. & Alcaraz, A. Single-molecule conformational dynamics of viroporin ion channels regulated by lipid-protein interactions. Bioelectrochemistry 137, 107641 (2021).

33. OuYang, B. & Chou, J.J. The minimalist architectures of viroporins and their therapeutic implications. Biochim Biophys Acta 1838, 1058–1067 (2014).

34. Miller, A.N. et al. The SARS-CoV-2 accessory protein Orf3a is not an ion channel, but does interact with trafficking proteins. Elife 12 (2023).

35. Qiu, F. & Dahl, G. A permeant regulating its permeation pore: inhibition of pannexin 1 channels by ATP. Am J Physiol Cell Physiol 296, C250–255 (2009).

36. Qiu, F., Wang, J. & Dahl, G. Alanine substitution scanning of pannexin1 reveals amino acid residues mediating ATP sensitivity. Purinergic Signal 8, 81–90 (2012).

37. Wang, J., Jackson, D.G. & Dahl, G. Cationic control of Panx1 channel function. Am J Physiol Cell Physiol. 315, C279–C289 (2018).

38. Michalski, K. & Kawate, T. Carbenoxolone inhibits Pannexin1 channels through interactions in the first extracellular loop. J Gen Physiol 147, 165–174 (2016).

39. Chekeni, F.B. et al. Pannexin 1 channels mediate ’find-me’ signal release and membrane permeability during apoptosis. Nature 467, 863–867 (2010).

40. Chiu, Y.H., Ravichandran, K.S. & Bayliss, D.A. Intrinsic properties and regulation of Pannexin 1 channel. Channels (Austin) 8, 1–7 (2014).

41. Silverman, W., Locovei, S. & Dahl, G. Probenecid, a gout remedy, inhibits pannexin 1 channels. Am J Physiol Cell Physiol 295, C761–767 (2008).

42. Wang, J. & Dahl, G. SCAM analysis of Panx1 suggests a peculiar pore structure. J Gen Physiol 136, 515–527 (2010).

43. Slavi, N. et al. Connexin 46 (cx46) gap junctions provide a pathway for the delivery of glutathione to the lens nucleus. J Biol Chem 289, 32694–32702 (2014).

44. Stridh, M.H., Tranberg, M., Weber, S.G., Blomstrand, F. & Sandberg, M. Stimulated efflux of amino acids and glutathione from cultured hippocampal slices by omission of extracellular calcium: likely involvement of connexin hemichannels. J Biol Chem 283, 10347–10356 (2008).

45. Tong, X. et al. Glutathione release through connexin hemichannels: Implications for chemical modification of pores permeable to large molecules. J Gen Physiol 146, 245–254 (2015).

46. Bunse, S. et al. Single cysteines in the extracellular and transmembrane regions modulate pannexin 1 channel function. J Membr Biol 244, 21–33 (2011).

47. Wang, J., Jackson, D.G. & Dahl, G. The food dye FD&C Blue No. 1 is a selective inhibitor of the ATP release channel Panx1. J Gen Physiol 141, 649–656 (2013).

48. Gordon, L.G. & Haydon, D.A. The unit conductance channel of alamethicin. Biochim Biophys Acta 255, 1014–1018 (1972).

49. Pressman, B.C. Ionophorous antibiotics as models for biological transport. Fed Proc 27, 1283–1288 (1968).

50. Drumm, M.L. et al. Chloride conductance expressed by delta F508 and other mutant CFTRs in Xenopus oocytes. Science 254, 1797–1799 (1991).

51. Cheng, S.H. et al. Defective intracellular transport and processing of CFTR is the molecular basis of most cystic fibrosis. Cell 63, 827–834 (1990).

52. Riordan, J.R. Assembly of functional CFTR chloride channels. Annu Rev Physiol 67, 701–718 (2005).

53. Sick, T.J., Feng, Z.C. & Rosenthal, M. Spatial stability of extracellular potassium ion and blood flow distribution in rat cerebral cortex after permanent middle cerebral artery occlusion. J Cereb Blood Flow Metab 18, 1114–1120 (1998).

54. Kuzuya, M. et al. Structures of human pannexin-1 in nanodiscs reveal gating mediated by dynamic movement of the N terminus and phospholipids. Sci Signal 15, eabg6941 (2022).

55. Pettersen, E.F. et al. UCSF Chimera--a visualization system for exploratory research and analysis. J Comput Chem 25, 1605–1612 (2004).

